# Sensory neuron dysfunction and hyperexcitability in dorsal root ganglia at disease onset in the SOD1G93A mouse model of ALS

**DOI:** 10.64898/2026.08.27.747263

**Authors:** Akira Nishiura, Soju Seki, Takuhiro Kobayashi, Yoshihiro Kitaoka, Sou Kawata, Toru Yamamoto, Kazuaki Miyagawa, Toshihiro Uchihashi, Yosuke Ohtake, Takahide Itokazu, Toshihide Yamashita, Masayuki Ono, Yusuke Yokota, Emiko Tanaka Isomura, Susumu Tanaka

## Abstract

Amyotrophic lateral sclerosis (ALS) is a progressive neurodegenerative disorder traditionally characterized by motor neuron degeneration, but emerging evidence indicates sensory system involvement. Despite reports of sensory abnormalities in some patients, the molecular and functional alterations in dorsal root ganglion (DRG) neurons remain insufficiently characterized. We investigated DRG pathology at disease onset in 12-week-old SOD1G93A mice using an integrated transcriptomic, morphological, and electrophysiological approach. RNA sequencing of lumbar DRG identified 35 differentially expressed genes, predominantly upregulated, enriched in oxidative stress-related and phagosome pathways. Comparative analysis with motor neuron transcriptomes revealed distinct gene expression profiles, indicating sensory neuron-specific molecular responses. Immunohistochemistry demonstrated reduced soma diameter in both A-and C-fiber DRG neurons. Nav channel colocalization increased for Nav1.7 in A fibers and Nav1.8 in both fiber types, whereas Nav1.6 was unchanged. Whole-cell patch-clamp recordings showed depolarized resting membrane potential, increased spike amplitude, and enhanced repetitive firing in A-fiber neurons, consistent with hyperexcitability, while C fibers showed no significant functional changes. These findings demonstrate early molecular, structural, and functional alterations in primary sensory neurons in ALS, supporting pathology beyond motor neurons and identifying sensory neuron excitability as a potential therapeutic target.

## Introduction

Amyotrophic lateral sclerosis (ALS) is a progressive neurodegenerative disorder characterized by the degeneration and loss of upper and lower motor neurons in the cerebral cortex, brainstem, and spinal cord. The incidence is approximately 2 per 100,000 individuals, with a typical age at onset of 50–60 years^[1]^. Clinically, ALS presents with progressive muscle weakness and atrophy, leading to limb paralysis and bulbar symptoms such as dysarthria and dysphagia, and ultimately resulting in death because of respiratory failure within 3–5 years of disease onset^1^. Currently approved therapeutic agents, including riluzole and edaravone, provide only modest disease-modifying effects, and no curative treatment is available^[2–4]^. A more comprehensive understanding of the underlying pathophysiological mechanisms is therefore essential for the development of effective therapeutic strategies.

Although ALS has traditionally been considered a disorder selectively affecting the motor system, increasing evidence indicates involvement of non-motor systems, including sensory and cognitive domains^[5]^. Sensory abnormalities have been reported in approximately 20%– 30% of patients, manifesting as distal limb pain, numbness, or hypoesthesia^[6]^. Despite these observations, pathological alterations in primary sensory neurons of the peripheral nervous system remain insufficiently characterized.

The cell bodies of primary sensory neurons reside in the dorsal root ganglia (DRG), which relay somatosensory information from the periphery to the central nervous system^[7]^. DRG neurons are broadly classified into Aβ, Aδ, and C fiber subtypes, which mediate mechanosensation, fast nociception, and slow nociception and temperature sensation, respectively^[8]^. Neuronal excitability in these cells is critically regulated by voltage-gated sodium (Nav) channels, and dysregulation of Nav channel expression or function is strongly implicated in neuropathic pain and other sensory disturbances^[9,10]^. Among these, Nav1.6, Nav1.7, and Nav1.8 are prominently expressed in DRG neurons and are susceptible to modulation by oxidative stress and inflammatory mediators^[9–12]^. These findings raise the possibility that molecular and electrophysiological alterations in DRG neurons may contribute to sensory dysfunction in ALS^[5,6]^.

Approximately 95% of ALS cases are sporadic, whereas the remaining 5% are familial, with mutations in the superoxide dismutase 1 (*SOD1*) gene representing one of the established genetic causes^[3,13]^. The *SOD1^G93A^* transgenic mouse model is widely used in ALS research and has been extensively characterized with respect to motor neuron pathology^[14,15]^. More recently, alterations in firing properties and Nav-dependent excitability have also been described in primary sensory neurons, including mesencephalic trigeminal neurons, in this model^[16,17]^. However, comprehensive analyses integrating molecular, morphological, and electrophysiological changes in limb DRG neurons remain limited.

The *SOD1^G93A^* mutation confers a toxic gain of function associated with increased production of reactive oxygen species, and oxidative stress is considered a major contributor to motor neuron degeneration^[18,19]^. In contrast, the impact of oxidative stress–related gene alterations in the DRG and their potential effects on sensory neuron excitability have not been fully elucidated.

In the present study, we aimed to investigate DRG neurons in SOD1^G93A^ mice at disease onset using an integrated approach combining RNA sequencing (RNA-seq), immunohistochemistry, and electrophysiological recordings. We performed comparative analyses with motor neurons to identify molecular alterations specific to primary sensory neurons and to determine whether ALS-associated pathology extends to the sensory system, with particular emphasis on Nav channel–mediated changes in membrane excitability.

## Results

### Gene expression analysis of DRG

RNA-seq was performed on lumbar DRG tissues obtained from 12-week-old WT and mSOD1 mice (Fig. 1A). To further examine transcriptional patterns underlying genotype separation, a heatmap was generated using the top 1,000 genes ranked by expression level. Non-hierarchical clustering (k = 3, determined by the elbow method) identified three clusters. Clusters 1 and 2 primarily reflected variation along PC1, whereas cluster 3 corresponded to genotype-associated separation along PC2 (Fig. 1B). Genes with the highest loading values on PC1 were predominantly housekeeping-related transcripts, including *Rps18-ps6* and *Eno1b*, whereas genes contributing strongly to PC2 included inflammation-and nerve injury– associated genes such as *Gfap*, *C3*, *Cxcl14*, and *Sprr1a* (Fig. 1B). PCA of all expressed genes (n = 47,709) revealed genotype-associated separation along the PC2 axis (PC1: 33.78% variance explained; PC2: 24.9%) (Fig. 1C).

**Fig. 1.**
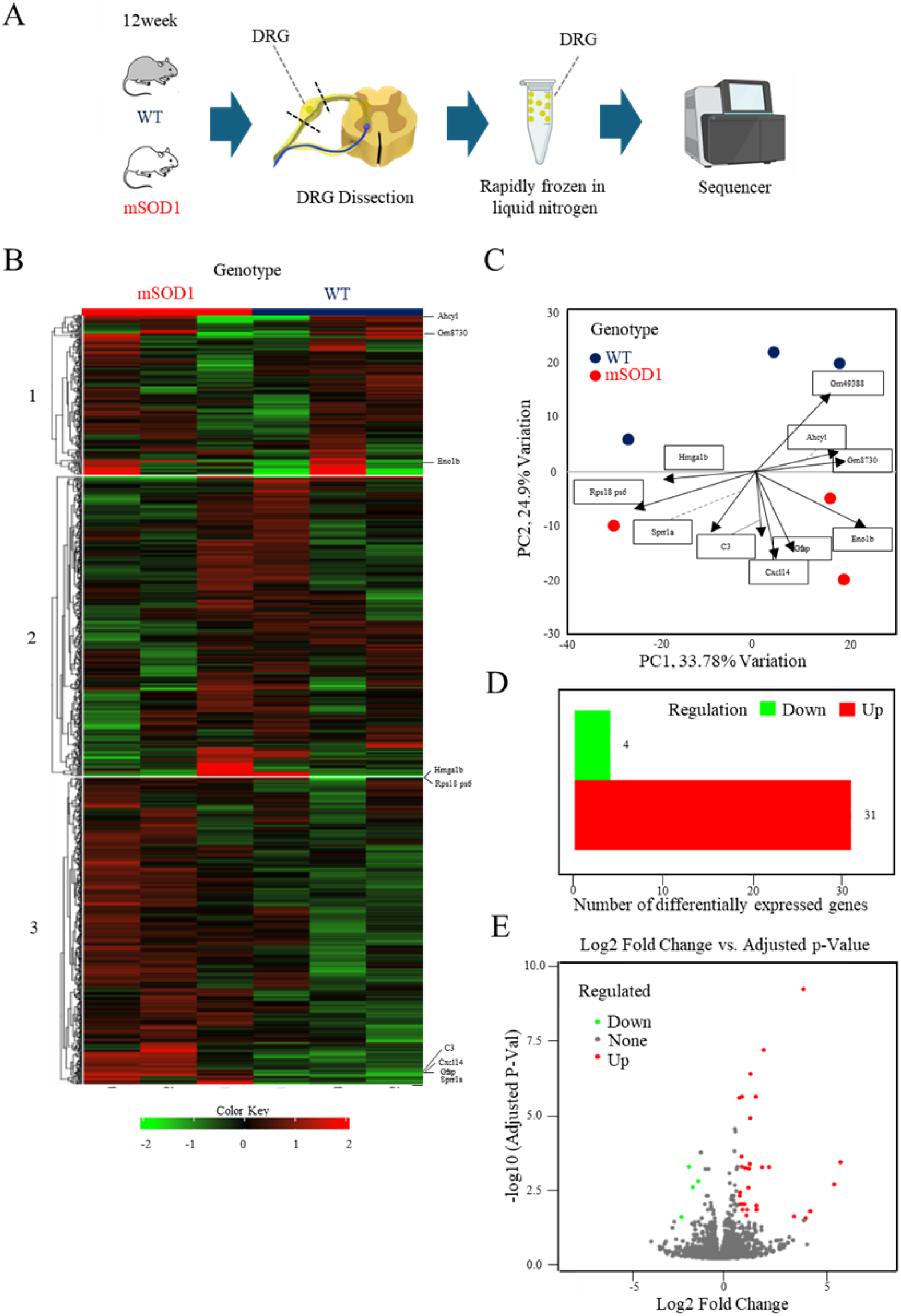
Gene expression profiling of DRG in mSOD1 mice. (A) Experimental design of the RNA-seq analysis of DRGs obtained from 12-week-old WT and mSOD1 mice. (B) Heatmap based on unsupervised clustering (k-means, k = 3) using the top 1,000 genes by expression level. Colors indicate normalized expression levels of each gene. (C) Principal component analysis (PCA). PCA was performed using RNA-seq data from DRG samples of WT and mSOD1 groups (total number of expressed genes: 47,709). Arrows indicate representative genes with high contributions to the principal components (WT: N = 3; mSOD1: N = 3). (D) Differential gene expression analysis. The genes satisfying a significance threshold of false discovery rate (FDR) < 0.05 and |fold change| > 2 were considered significantly differentially expressed genes. (E) Volcano plot. The x-axis indicates log_2_ fold change (mSOD1/WT), and the y-axis indicates −log_10_(adjusted *p*-value). Upregulated genes are shown in red and downregulated genes in green.

Differential expression analysis identified 35 significantly altered genes (FDR < 0.05, |fold change| > 2), of which 31 were upregulated and 4 were downregulated in mSOD1 mice compared with WT mice (Tables 1 and 2; Fig. 1D). A volcano plot illustrated the predominance of upregulated transcripts in the mSOD1 group (Fig. 1E). Furthermore, GSEA using the differentially expressed genes (DEGs) demonstrated significant enrichment of the phagosome pathway (adjusted *p* = 3.8 × 10⁻⁴) (Table 3; Fig. 2A). Mapping DEGs onto the KEGG phagosome map indicated upregulation of multiple genes, involved in phagosomal function, including those involved in NADPH oxidase complex formation, phagosome maturation, and reactive oxygen species (ROS) production, were upregulated in the mSOD1 group (Table 4; Fig. 2B).

**Fig. 2.**
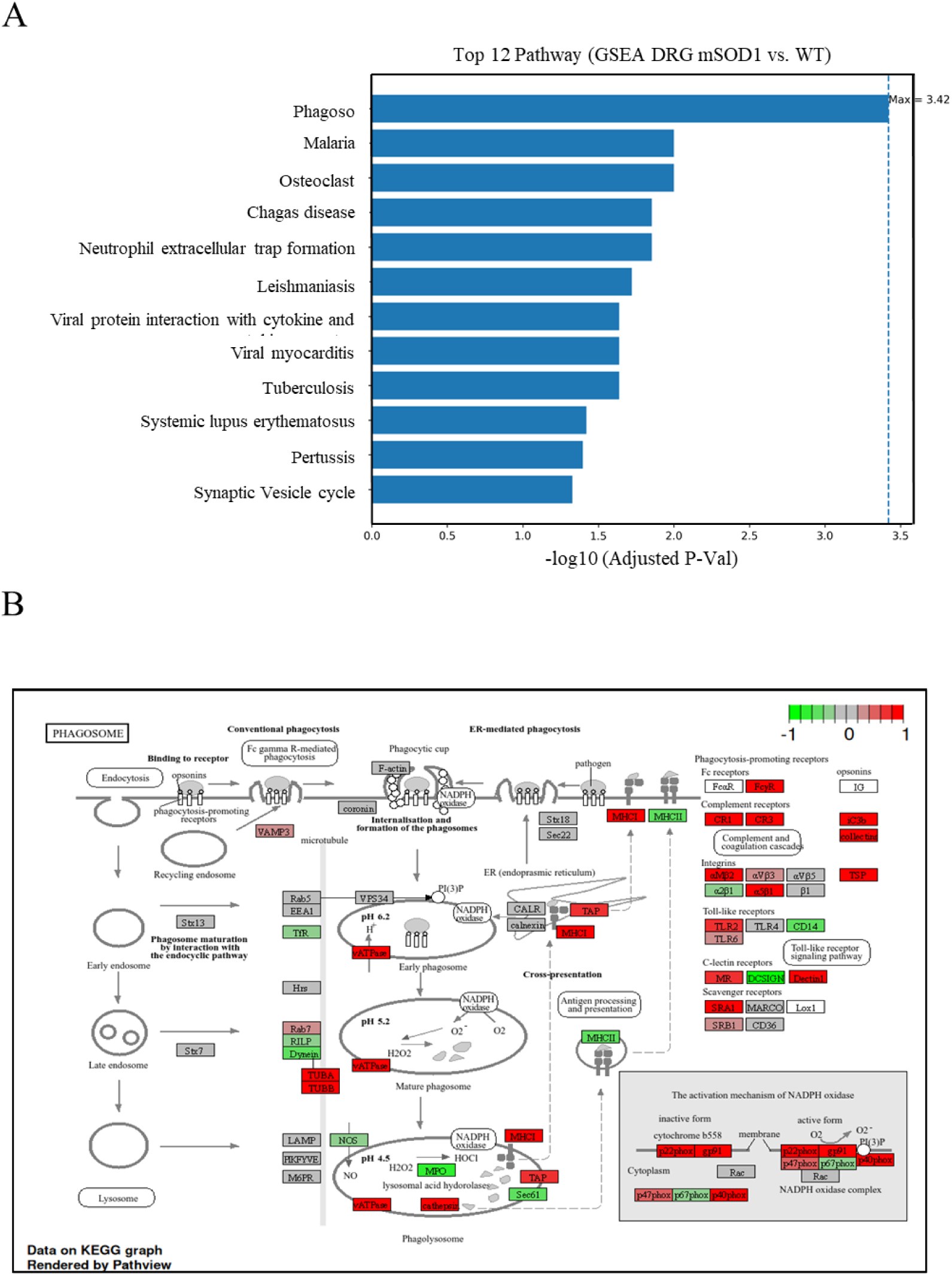
Functional pathway analysis in DRG of mSOD1 mice. (A) Pathway analysis by Gene Set Enrichment Analysis (GSEA). GSEA was performed using the set of differentially expressed genes (DEGs), and the top 12 pathways with the highest significance are shown. The x-axis indicates −log_10_ (adjusted *p*-value). (B) KEGG phagosome pathway map. Red indicates upregulated genes, and green indicates downregulated genes.

**Table 1.**
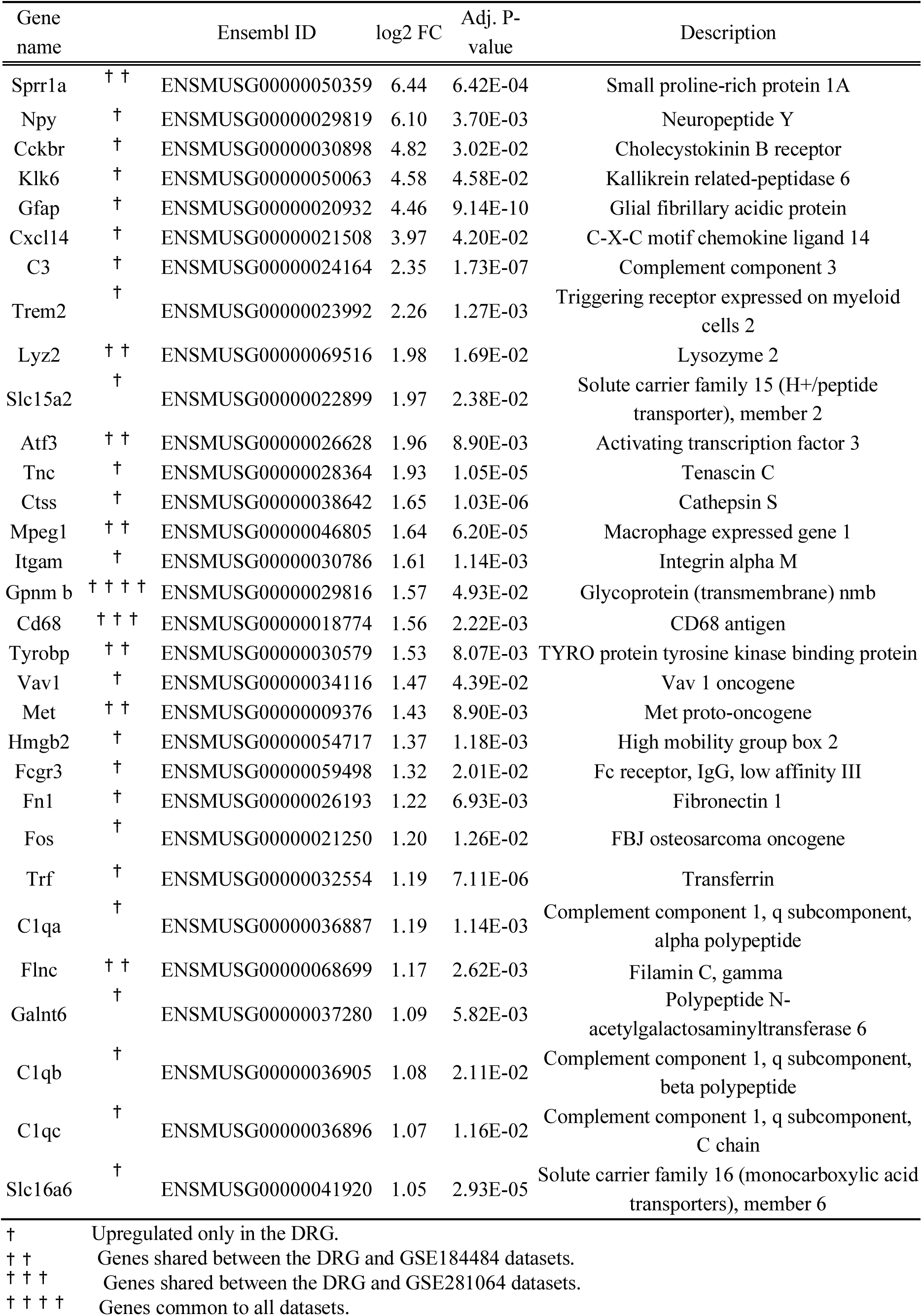
Genes upregulated in the DRG.

**Table 2.** Genes downregulated in the DRG.

| Gene name | Ensembl ID | log2 FC | Adj. P-value | Description |
| --- | --- | --- | --- | --- |
| Dlk1 | ENSMUSG00000040856 | -2.03 | 4.42E-02 | Delta like non-canonical Notch ligand 1 |
| Ednra | ENSMUSG00000031616 | -1.63 | 1.11E-03 | Endothelin receptor type A |
| Plk5 | ENSMUSG00000035486 | -1.42 | 1.55E-03 | Polo like kinase 5 |
| Serpind1 | ENSMUSG00000022766 | -1.13 | 4.15E-03 | Serine (or cysteine) peptidase inhibitor, clade D, member 1 |

**Table 3.** Pathway analysis by GSEA.

| Rank | Pathway | NES | Adj. P-value |
| --- | --- | --- | --- |
| 1 | Phagosome | 0.62 | 3.80E-04 |
| 2 | Malaria | 0.74 | 1.00E-02 |
| 3 | Osteoclast differentiation | 0.57 | 1.00E-02 |
| 4 | Chagas disease | 0.59 | 1.40E-02 |
| 5 | Neutrophil extracellular trap formation | 0.57 | 1.40E-02 |
| 6 | Leishmaniasis | 0.66 | 1.90E-02 |
| 7 | Viral protein interaction with cytokine and cytokine receptor | 0.71 | 2.30E-02 |
| 8 | Viral myocarditis | 0.63 | 2.30E-02 |
| 9 | Tuberculosis | 0.52 | 2.30E-02 |
| 10 | Systemic lupus erythematosus | 0.66 | 3.80E-02 |
| 11 | Pertussis | 0.61 | 4.00E-02 |
| 12 | Synaptic vesicle cycle | 0.61 | 4.70E-02 |

**Table 4.**
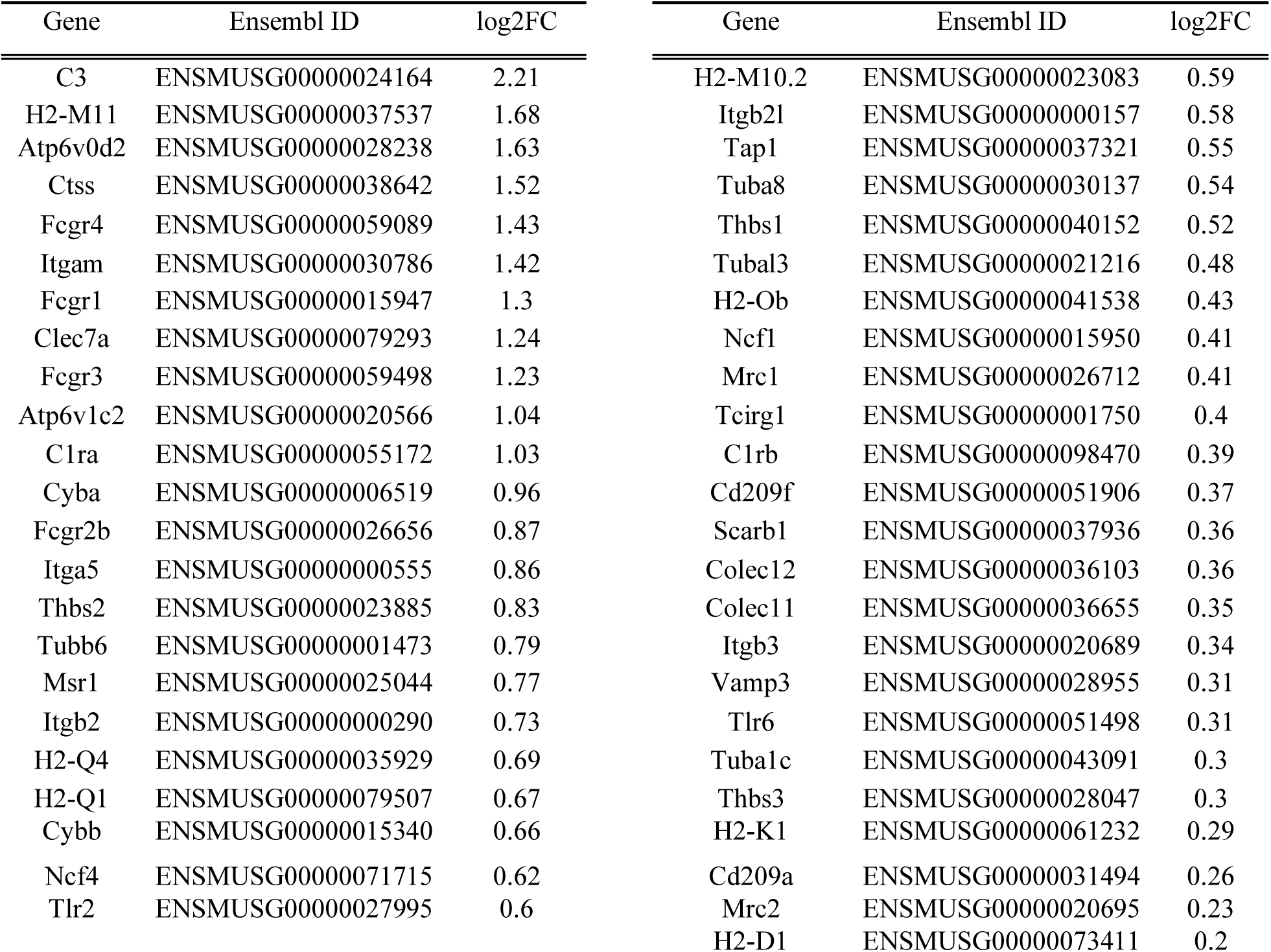
Genes Altered in the Phagosome Pathway.

### Comparative analysis between DRG and spinal motor neurons

Comparison of our dataset with publicly available motor neuron (MN) transcriptomes (GSE184484 and GSE281064) identified only one gene, *Gpnmb*, consistently increased expression across all three datasets. In contrast, 22 genes were selectively upregulated in mSOD1 DRG, including *Fos* and *Neuropeptide Y (Npy)*.

Eight genes were commonly upregulated across both MN datasets, including ALS-related genes such as *Cd109*, *Lgals3*, and *Timp1*. When overlaps were analyzed separately, seven genes were shared between DRG and GSE184484, including inflammation-and immune-related genes (*Tyrobp*, *Mpeg1*, *Lyz2*) and injury-response genes (*Sprr1a*, *Met*). Only one gene, *Cd68*, overlapped between DRG and GSE281064 (Fig. 3). In contrast, 72 genes were uniquely upregulated in GSE184484 and 15 genes in GSE281064 (Fig. 3).

**Fig. 3.**
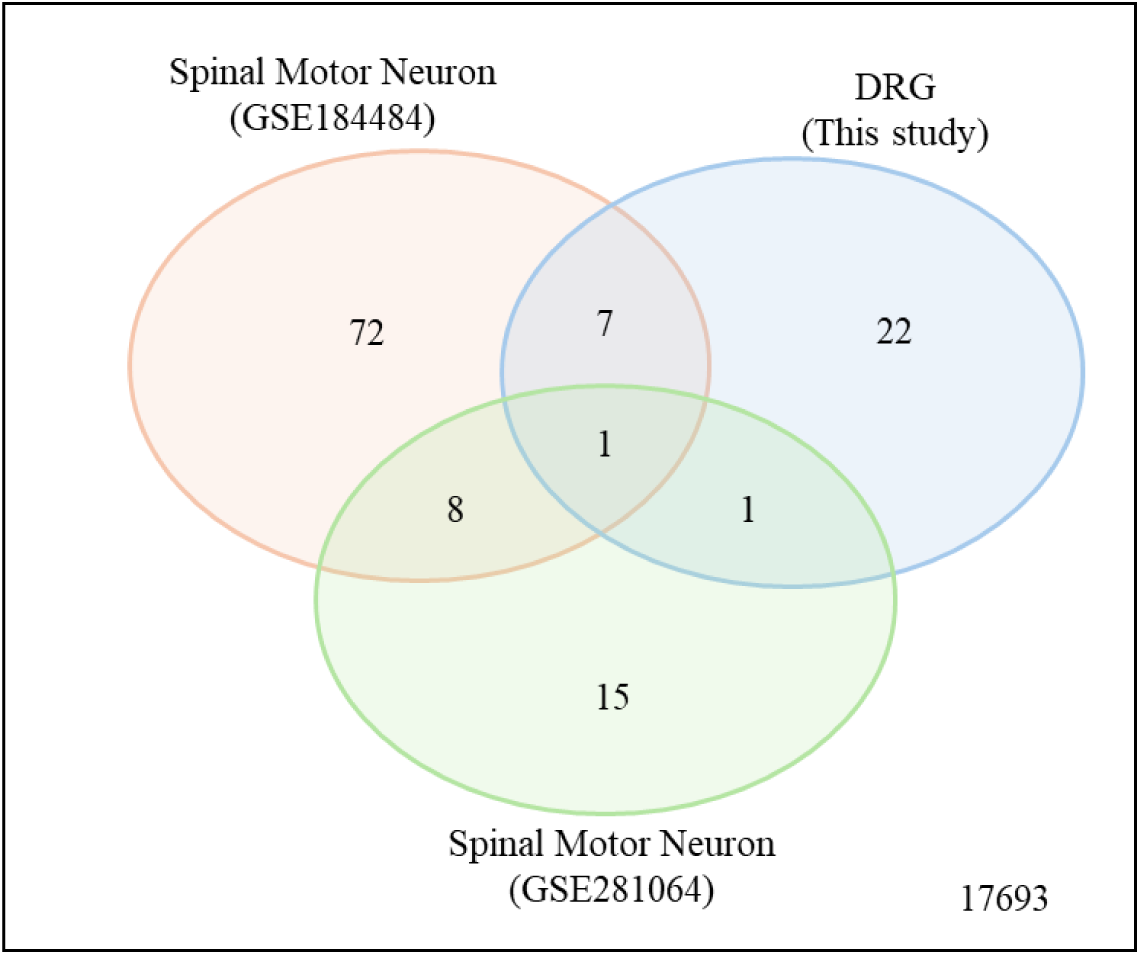
Comparative analysis between DRG and MNs in mSOD1 mice. Comparative analysis of RNA-seq results from mSOD1 mouse DRG (this study) and publicly available RNA-seq datasets of MNs from age-matched mSOD1 mice (GSE184484 and GSE281064). The overlap of upregulated genes identified by differential gene expression analysis in each dataset is shown.

### Quantitative analysis of neuronal soma size in DRG neurons

Fluorescence immunostaining was performed on DRG sections from WT and mSOD1 mice using NF200 (A-fiber marker) and peripherin (C-fiber marker), and the long-axis diameter of neuronal somata was measured (Fig. 4A, B). In A fibers, the long-axis diameter was significantly reduced in mSOD1 mice compared with that in WT mice (WT: 32.77 ± 4.64 µm, n = 53; mSOD1: 27.95 ± 5.44 µm, n = 56; Welch’s t-test, *p* < 0.001) (Fig. 4C). Similarly, C fibers exhibited a significant reduction in soma diameter in mSOD1 group (23.0 ± 5.44 µm, n = 57) compared with that in WT group (26.41 ± 5.64 µm, n = 62) (*p* = 0.001; Fig. 4D).

**Fig. 4.**
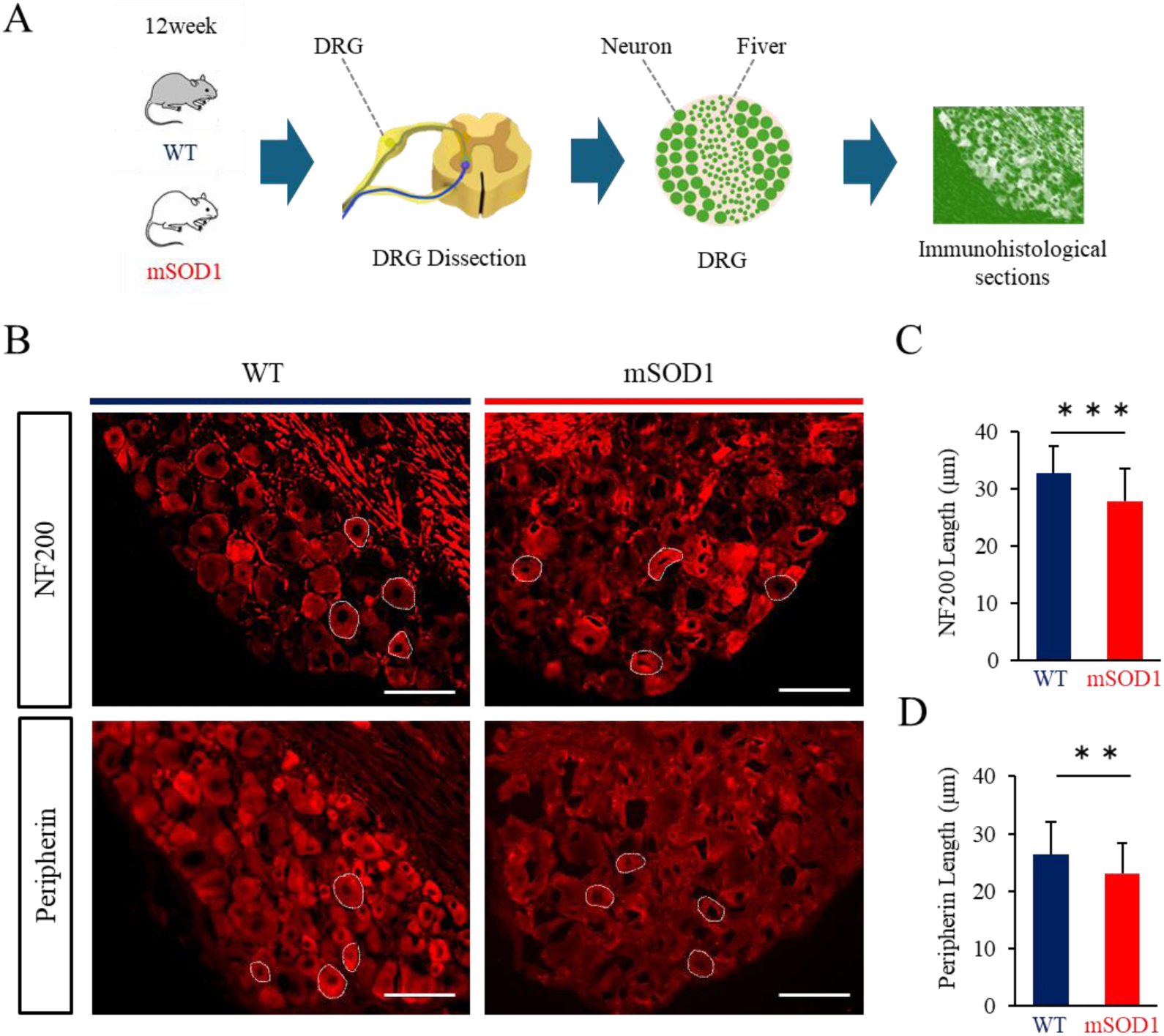
Quantitative analysis of DRG neuron soma diameter. (A) Experimental design of the immunohistochemical analysis of DRGs obtained from 2-week-old WT and mSOD1 mice. (B) Immunofluorescence images of NF200-positive neurons (upper panels) and peripherin-positive neurons (lower panels) in the DRG of WT and mSOD1 mice. NF200 indicates A-fiber–derived neurons, and peripherin indicates C-fiber–derived neurons. The regions outlined in white indicate the regions of interest (ROIs) set for measuring the long diameter of the neuronal soma (scale bar: 50 μm). (C) Quantification of the soma long diameter of NF200-positive neurons; WT: n = 53, mSOD1: n = 56, N = 3 per group, mean ± SD, Welch’s t-test, *p* < 0.001, *** *p* < 0.001. (D) Quantification of the soma long diameter of Peripherin-positive neurons. WT: n = 62, mSOD1: n = 57, N = 3 per group, mean ± SD, Welch’s t-test, *p* = 0.001, ** *p* < 0.01.

### Analysis of Nav channel colocalization in DRG neurons

Double immunofluorescence staining for NF200 (A-fiber marker) or peripherin (C-fiber marker) together with Nav1.6, Nav1.7, or Nav1.8 was performed on DRG sections from WT and mSOD1 mice. Colocalization ratios of Nav channels within identified neuronal subtypes were quantified.

For Nav1.6, no significant differences in colocalization ratios were observed between groups in either A fibers (WT: n = 9, 0.43 ± 0.06; mSOD1: n = 9, 0.47 ± 0.04; Welch’s t-test, *p* = 0.134) or C fibers (WT: n = 9, 0.42 ± 0.11; mSOD1: n = 9, 0.49 ± 0.12; *p* = 0.214) (Fig. 5A– C).

**Fig. 5.**
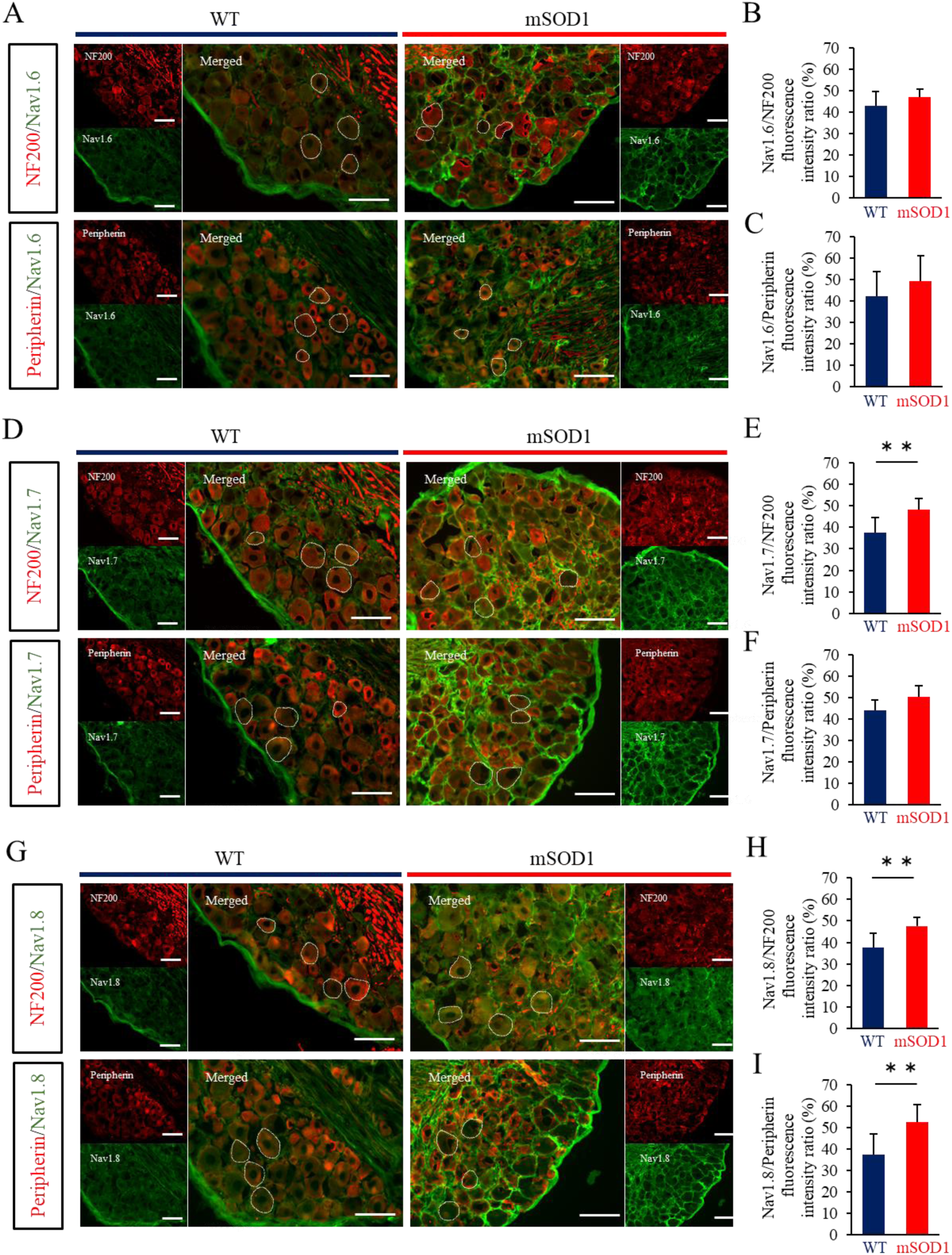
Colocalization ratios of Nav1.6, Nav1.7, and Nav1.8 in DRG neurons. (A) Double immunofluorescence images of Nav1.6 with NF200 (upper panels) or peripherin (lower panels) in the DRG of WT and mSOD1 mice. NF200 indicates A-fiber–derived neurons, and peripherin indicates C-fiber–derived neurons. Merged images show the overlap of each Nav channel (green) with the fiber markers (red). Regions outlined in white indicate the regions of interest (ROIs) set for quantitative analysis (scale bar: 50 μm). (B) Quantification of the Nav/NF200 fluorescence intensity ratio in NF200-positive neurons. WT: n = 9, 0.43 ± 0.06; mSOD1: n = 9, 0.47 ± 0.04; Welch’s t-test, *p* = 0.134. (C) Quantification of the Nav/peripherin fluorescence intensity ratio in peripherin-positive neurons. WT: n = 9, 0.42 ± 0.11; mSOD1: n = 9, 0.49 ± 0.12; Welch’s t-test, *p* = 0.214. (D) Double immunofluorescence images of Nav1.7 with NF200 (upper panels) or peripherin (lower panels). (E) Quantification of the Nav/NF200 fluorescence intensity ratio in NF200-positive neurons. WT: n = 9, 0.38 ± 0.07; mSOD1: n = 9, 0.48 ± 0.05; Welch’s t-test, *p* = 0.002, ** *p* < 0.01. (F) Quantification of the Nav/peripherin fluorescence intensity ratio in peripherin-positive neurons. WT: n = 9, 0.49 ± 0.07; mSOD1: n = 9, 0.50 ± 0.14; Welch’s t-test, *p* = 0.230. (G) Double immunofluorescence images of Nav1.8 with NF200 (upper panels) or peripherin (lower panels). (H) Quantification of the Nav/NF200 fluorescence intensity ratio in NF200-positive neurons. WT: n = 9, 0.38 ± 0.07; mSOD1: n = 9, 0.48 ± 0.04; Welch’s t-test, *p* = 0.002, ** *p* < 0.01. (I) Quantification of the Nav/peripherin fluorescence intensity ratio in Peripherin-positive neurons. WT: n = 9, 0.37 ± 0.10; mSOD1: n = 9, 0.53 ± 0.08; Welch’s t-test, *p* = 0.002, ** *p* < 0.01.

For Nav1.7, the colocalization ratio was significantly increased in A fibers of mSOD1 mice compared with that in WT mice (WT: n = 9, 0.38 ± 0.07; mSOD1: n = 9, 0.48 ± 0.05; Welch’s t-test, *p* = 0.002), whereas no significant difference was observed in C fibers (WT: n = 9, 0.49 ± 0.07; mSOD1: n = 9, 0.50 ± 0.14; Welch’s t-test, *p* = 0.230) (Fig. 5D–F).

For Nav1.8, colocalization ratios were significantly increased in mSOD1 mice in both A fibers (WT: n = 9, 0.38 ± 0.07; mSOD1: n = 9, 0.48 ± 0.04; Welch’s t-test, *p* = 0.002) and C fibers (WT: n = 9, 0.37 ± 0.10; mSOD1: n = 9, 0.53 ± 0.08; Welch’s t-test, *p* = 0.002) (Fig. 5G– I).

### Electrophysiological changes in DRG neurons

Based on the increased Nav channel colocalization observed in mSOD1 mice, electrophysiological properties of DRG neurons were examined (Fig. 6A). Neurons were classified as A fibers or C fibers according to their voltage responses to hyperpolarizing current pulses under current-clamp conditions (Fig. 6B) ^[20]^. In addition, neurons were categorized into two groups according to their responses to rectangular depolarizing current pulses: those exhibiting a single spike and those exhibiting repetitive firing (Fig. 6C).

**Fig. 6.**
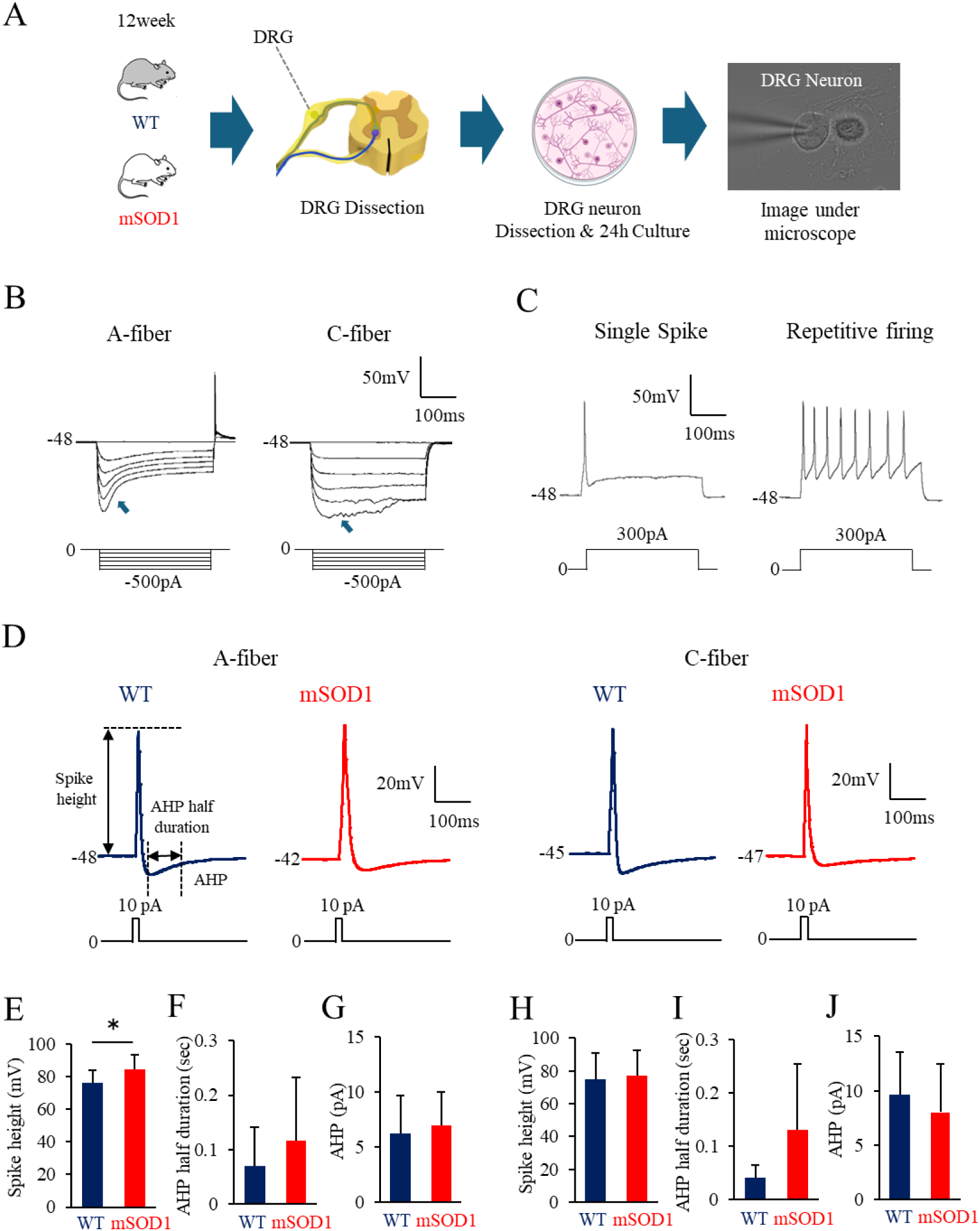
Whole-cell recordings of DRG neurons in mSOD1 mice. (A) Experimental design of the electrophysiological analysis of DRG neurons obtained from 12-week-old WT and mSOD1 mice. (B) A-fiber and C-fiber neurons were classified based on the presence or absence of a voltage sag (arrow) in the membrane voltage response during current-clamp recordings with hyperpolarizing current pulses. (C) Whole-cell recordings of DRG neurons in the WT and mSOD1 groups. Representative traces showing a single spike and repetitive firing evoked by a 0.5 s current injection of 200 pA are presented (WT: n = 20, mSOD1: n = 26). (D) In DRG neurons from the WT and mSOD1 groups, action potential (AP) properties were analyzed under current-clamp conditions by applying a rectangular depolarizing current pulse (10 pA, 10 ms). Representative AP waveforms from A-fiber–derived DRG neurons and C-fiber–derived DRG neurons in the WT and mSOD1 groups are shown. (E-G) These graphs present the quantification of spike height, AHP half duration, and AHP. WT = 7, mSOD1 = 12, mean ± SD, Welch’s t-test, spike height: *p* = 0.027, * *p* < 0.05. (H-J) In C-fiber–derived DRG neurons, no significant differences were observed between the WT and mSOD1 groups in any of the measured parameters, including spike height, AHP half duration, and AHP (WT = 7, mSOD1 = 10, Welch’s t-test, spike height: *p* = 0.789; AHP half duration: *p* = 0.104; AHP: *p* = 0.504).

Regarding basic membrane properties, A fibers in the mSOD1 group exhibited a significantly depolarized RMP compared with WT controls (WT: n = 13, −48.11 ± 6.63 mV; mSOD1: n = 13, −42.43 ± 4.76 mV; Welch’s t-test, *p* = 0.046). No significant differences were observed in Cm or Rin (Table 5). In C fibers, no significant differences were observed in RMP, Cm, or Rin (Table 5).

**Table 5.**
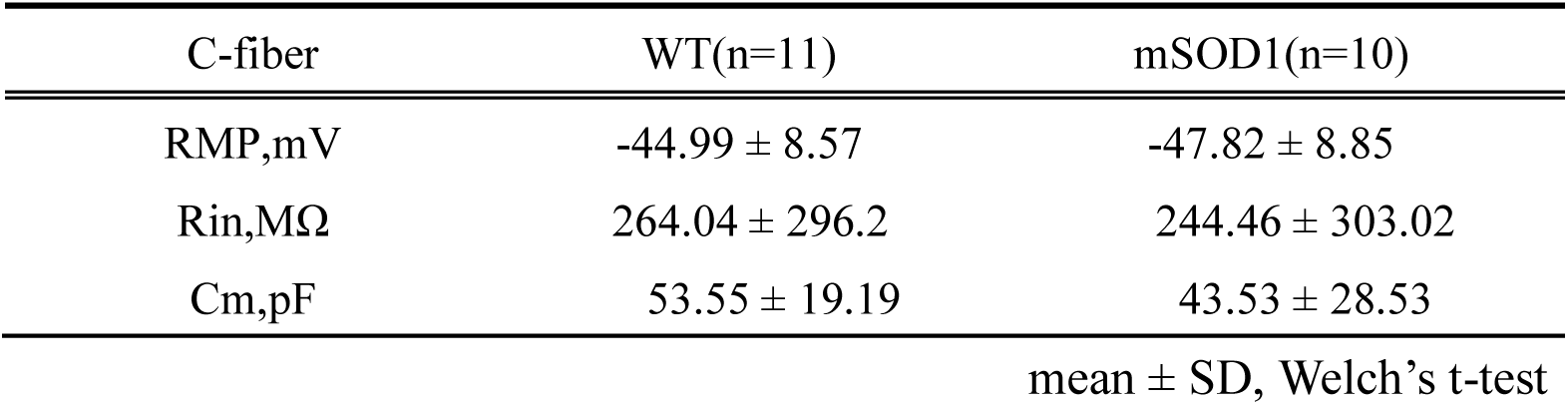
Basic membrane properties.

| C-fiber | WT(n=11) | mSOD1(n=10) |
| --- | --- | --- |
| RMP,mV | -44.99 ± 8.57 | -47.82 ± 8.85 |
| Rin,MΩ | 264.04 ± 296.2 | 244.46 ± 303.02 |
| Cm,pF | 53.55 ± 19.19 | 43.53 ± 28.53 |

mean ± SD, Welch's t-test

Under current-clamp conditions, depolarizing pulses (10 pA, 10 ms) evoked APs (Fig. 6D). In A fibers, spike amplitude was significantly increased in mSOD1 mice (WT: n = 7, 76.42 ± 7.25 mV; mSOD1: n = 12, 84.33 ± 9.14 mV; *p* = 0.027) (Fig. 6E). No significant differences were detected in AHP amplitude or AHP half-duration (AHP: WT: n = 7, 6.26 ± 3.43 mV; mSOD1: n = 12, 6.94 ± 3.07 mV; AHP half duration: WT: n = 7, 0.07 ± 0.07 s; mSOD1: n = 12, 0.12 ± 0.12 s; Welch’s t-test, AHP: p = 0.271, AHP half duration: *p* = 0.191) (Table 6; Fig. 6F,G). In C fibers, no significant differences were observed in any AP parameters, including spike height, AHP, or AHP half duration (WT: n = 7, mSOD1: n = 10; spike height: WT: 74.95 ± 15.84 mV; mSOD1: 77.05 ± 15.3 mV; AHP: WT: 9.61 ± 3.96 mV; mSOD1: 8.1 ± 4.42 mV; AHP half duration: WT: 0.04 ± 0.02 s; mSOD1: 0.13 ± 0.12 s; Welch’s t-test, spike height: *p* = 0.789, AHP: *p* = 0.504, AHP half duration: *p* = 0.104) (Table 6; Fig. 6H-J).

**Table 6.** Action potential properties.

| A-fiber | WT(n=7) | mSOD1(n=12) | 832 |
| --- | --- | --- | --- |
| AP |  |  |  |
| Spike Height,mV | 76.42 ± 7.25 | 84.31±9.14 * |  |
| AHP |  |  |  |
| AHP peak,mV | 6.26 ± 3.43 | 6.94 ± 3.07 |  |
| Half duration,s | 0.07 ± 0.07 | 0.12 ± 0.12 |  |

mean ± SD, Welch's t-test

Application of step current injections (0–600 pA) distinguished neurons exhibiting single versus repetitive firing (Fig. 7A). In A fibers, rheobase showed a decreasing trend in mSOD1 group (WT: n = 3, 200 [200–300] pA; mSOD1: n = 8, 200 [150–200] pA; Mann–Whitney U test, *p* = 0.085) (Fig. 7B). In C fibers, no significant difference in rheobase was observed between groups (WT: n = 6, 400 (300–500) pA; mSOD1: n = 3, 400 (250–650) pA; Mann– Whitney’s U-test, *p* = 0.952) (Fig. 7C).

**Fig. 7.**
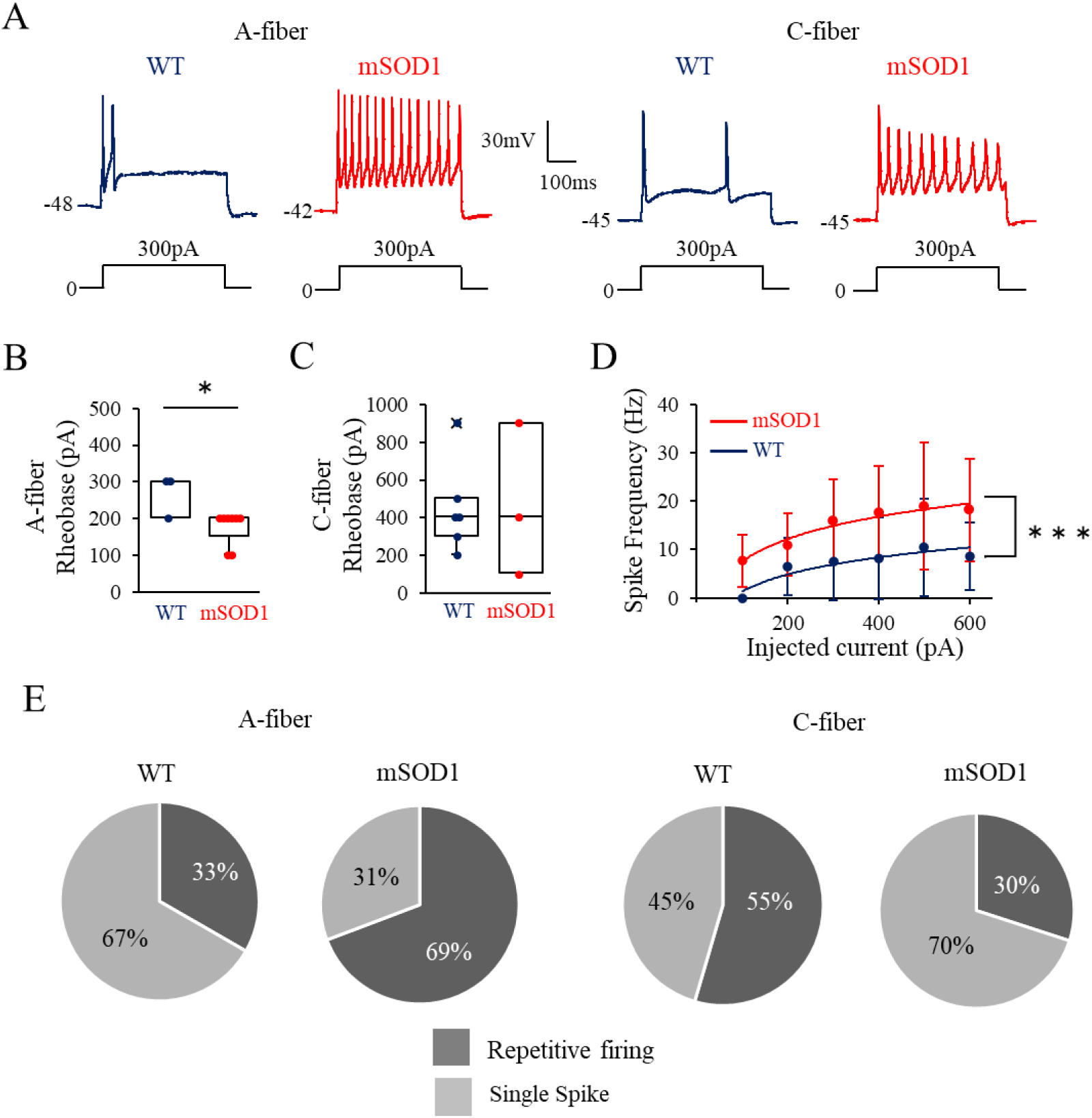
Repetitive firing properties of DRG neurons in mSOD1 mice. (A) Representative traces of repetitive firing in DRG neurons derived from each fiber type in the WT and mSOD1 groups in response to a hyperpolarizing pulse (300 pA, 0.5 s). (B) Quantification of rheobase in A-fiber–derived DRG neurons. Each dot represents an individual neuron, and the horizontal line indicates the median. WT: n = 3, 300 (200–300) pA; mSOD1: n = 8, 200 (150–200) pA; Mann–Whitney U test, *p* = 0.085, § *p* < 0.1. (C) Quantification of rheobase in C-fiber–derived DRG neurons. WT: n = 6, 400 (300–500) pA; mSOD1: n = 3, 400 (250–650) pA; Mann–Whitney U test, *p* = 0.952. (D) Comparison of frequency–current (F–I) curves in DRG neurons from WT and mSOD1 groups. Each plot indicates the mean firing frequency at each injected current. 200 pA: WT: n = 5, 6.58 ± 2.66 Hz; mSOD1: n = 10, 11.03 ± 2.05 Hz; 300 pA: WT: n = 6, 7.64 ± 3.31 Hz; mSOD1: n = 9, 15.87 ± 2.87 Hz; 400 pA: WT: n = 7, 8.21 ± 3.17 Hz; mSOD1: n = 9, 17.68 ± 3.19 Hz; 500 pA: WT: n = 7, 10.49 ± 3.81 Hz; mSOD1: n = 4, 18.96 ± 6.58 Hz; 600 pA: WT: n = 6, 8.65 ± 2.88 Hz; mSOD1: n = 3, 18.18 ± 6.07 Hz; two-way ANOVA, *p* = 0.0009, *** *p* < 0.001. (E) Proportion of repetitive firing in DRG neurons from WT and mSOD1 groups. For A-fiber–derived neurons (upper panels) and C-fiber–derived neurons (lower panels), the proportions of cells exhibiting single spikes and repetitive firing in response to hyperpolarizing pulses were compared. WT: repetitive firing n = 3, single spike n = 6; mSOD1: repetitive firing n = 9, single spike n = 4; Pearson’s chi-square test, *p* = 0.099. In contrast, no significant difference in firing proportions was observed between WT and mSOD1 groups in C-fiber–derived neurons. WT: repetitive firing n = 6, single spike n = 5; mSOD1: repetitive firing n = 3, single spike n = 7; Pearson’s chi-square test, *p* = 0.256.

Analysis of F–I curves in neurons exhibiting repetitive firing demonstrated a significant increase in mean firing frequency in mSOD1 mice compared with WT mice (200 pA: WT: n = 5, 6.58 ± 2.66 Hz; mSOD1: n = 10, 11.03 ± 2.05 Hz; 300 pA: WT: n = 6, 7.64 ± 3.31 Hz; mSOD1: n = 9, 15.87 ± 2.87 Hz; 400 pA: WT: n = 7, 8.21 ± 3.17 Hz; mSOD1: n = 9, 17.68 ± 3.19 Hz; 500 pA: WT: n = 7, 10.49 ± 3.81 Hz; mSOD1: n = 4, 18.96 ± 6.58 Hz; 600 pA: WT: n = 6, 8.65 ± 2.88 Hz; mSOD1: n = 3, 18.18 ± 6.07 Hz; two-way ANOVA, *p* = 0.0009) (Fig. 7D).

In A fibers, the proportion of neurons exhibiting repetitive firing was higher in mSOD1 mice (WT: 3/9, 0.33; mSOD1: 9/13, 0.69), showing a trend toward significance (Pearson’s χ² test, *p* = 0.099; Fig. 7E). In C fibers, no significant difference in the proportion of repetitive firing neurons was observed between groups (*p* = 0.256) (Fig. 7E).

## Discussion

In this study, RNA-seq analysis of DRG from SOD1G93A mice at disease onset revealed activation of oxidative stress-related pathways, accompanied by increased expression of activity-associated genes, including *Fos* and *Npy*. Immunohistochemical analyses demonstrated soma atrophy in A-fiber DRG neurons and altered localization of Nav1.7 and Nav1.8. Electrophysiological recordings further indicated enhanced membrane excitability, characterized by RMP depolarization, increased spike height, and a reduced threshold for repetitive firing. Collectively, these findings demonstrate that ALS-associated pathology extends beyond MN and involves primary sensory neurons at molecular, structural, and functional levels. The observed hyperexcitability of A-fiber DRG neurons suggests a potential mechanistic basis for the sensory disturbances reported clinically.

RNA-seq analysis identified the phagosome pathway as the most significantly altered pathway in the DRG of SOD1G93A mice at disease onset. The phagosome pathway is closely linked to oxidative stress-related processes, not only through its role in autophagy and pathogen clearance but also via ROS generation mediated by NADPH oxidase^[8]^. Consistent with this, KEGG pathway analysis indicated upregulation of gene groups involved in ROS production and processing, suggesting an enhanced oxidative stress environment in the DRG of ALS model mice at disease onset.

The SOD1G93A mutation impairs the physiological function of Cu/Zn SOD1 in scavenging superoxide anions and confers a toxic gain-of-function that promotes excessive ROS accumulation. This dual effect disrupts redox homeostasis, contributes to mitochondrial dysfunction, and increases intracellular oxidative stress^[18,19]^. Previous studies have primarily focused on oxidative stress-induced neurodegeneration in spinal motor neurons as a driver of ALS onset and progression^[19]^. In contrast, our findings indicate that oxidative stress–related pathways are also activated in the DRG, where primary sensory neuron somata reside, at disease onset. These results suggest that the DRG is exposed to a pathological molecular environment characteristic of ALS from early stages. Such oxidative stress-associated alterations may contribute to the Nav channel redistribution and membrane hyperexcitability observed in primary sensory neurons in this study.

Comparison of transcriptomic profiles between MNs and DRG revealed both shared and sensory neuron–specific pathological responses in SOD1G93A mice at disease onset. Among upregulated genes, *Gpnmb* was the only transcript consistently increased in both DRG and MN datasets. *Gpnmb* encodes a neuroprotective protein implicated in ALS. In MNs, mutant SOD1G93A has been shown to interfere with *Gpnmb* maturation by disrupting glycosylation, thereby reducing protein stability and promoting ubiquitination and degradation. This impairment limits its protective capacity and may render MNs more vulnerable^[21]^. This type of an interaction has not been clearly demonstrated for WT SOD1, suggesting that it represents a mutation-specific pathological mechanism. Elevated *Gpnmb* levels have also been reported in the spinal cord of ALS model mice at disease onset and in the serum of patients with ALS^[21]^, supporting the notion that certain stress-response pathways are shared between motor and sensory neuronal compartments.

In contrast, 22 genes were selectively upregulated in the DRG, indicating sensory neuron– specific molecular responses. Among these, *Fos* and *Npy* are of particular interest because of their established roles in neuronal activity and injury responses. *Fos* is a well-recognized marker of neuronal activation, and its increased expression in the DRG suggests heightened neuronal activity or stress exposure. The present study demonstrated a significant reduction in DRG soma diameter at disease onset, consistent with early neuronal stress or injury. Peripheral nerve injury is known to induce *Fos* expression in DRG neurons, and *Fos* activation can, in turn, drive *Npy* expression in injured sensory neurons^[22,23]^. *Npy* is typically expressed at low levels in healthy DRG neurons but is markedly upregulated following peripheral nerve injury. It has been implicated in modulating excitability, particularly in A-fiber–derived DRG neurons^[23,24]^, and is considered a marker of A-fiber injury^[25]^. Increased *Npy* expression has also been associated with enhanced membrane excitability in other primary sensory neuron populations. Together, these findings suggest that selective upregulation of *Npy* in the DRG may contribute to enhanced excitability of A-fiber sensory neurons in SOD1G93A mice at disease onset.

RNA-seq analysis of DRG tissue from SOD1G93A mice at disease onset revealed coordinated changes in gene clusters associated with oxidative stress, complement activation, and immune responses. Comprehensive molecular characterization of the DRG at early disease stages has been limited in previous studies. Our findings suggest that, rather than reflecting activation of isolated trigger genes, the DRG exhibits a broader stress-responsive molecular signature, likely driven by mutation-induced oxidative imbalance and immune activation. In contrast to spinal MNs, where mitochondrial dysfunction, impaired proteostasis, and activation of apoptosis-related pathways have been reported from presymptomatic stages^[26,27]^, we did not detect clear upregulation of apoptosis-associated genes in the DRG at disease onset. This difference may indicate distinct vulnerability profiles or temporal dynamics between motor and sensory neuronal populations. Morphological analysis further demonstrated a significant reduction in DRG soma diameter in SOD1G93A mice, indicating that structural alterations are already present at disease onset. Previous studies have shown that cultured DRG neurons from ALS model mice exhibit impaired neurite outgrowth within 24 h^[28]^, suggesting intrinsic cellular vulnerability. The soma atrophy observed here may reflect a related early degenerative or stress-induced process. The oxidative stress-related transcriptional alterations may influence Nav channel expression and function, thereby contributing to changes in membrane excitability^[11,12,29]^. The molecular signatures identified in the DRG are therefore consistent with, and may mechanistically underlie, the electrophysiological alterations observed in this study.

Immunohistochemical analysis demonstrated significantly increased colocalization of Nav1.7 and Nav1.8 in A-fiber DRG neurons of ALS model mice at disease onset. In C fibers, Nav1.7 localization was unchanged, whereas Nav1.8 colocalization was increased, indicating fiber-type–specific alterations in Nav channel distribution. Nav1.7 and Nav1.8 are key determinants of excitability in primary sensory neurons^[9,10]^, contributing to action potential initiation and repetitive firing. In peripheral nerve injury models, increased expression of Nav1.7 and Nav1.8 has been consistently reported in DRG neurons^[30]^, supporting their role in pathological hyperexcitability. The Nav channel redistribution observed in the DRG at ALS disease onset may therefore represent a conserved molecular response to neuronal stress or injury. Alterations in Nav channel expression and localization are common features of neuropathic conditions and are considered central to the development of sensory neuron dysfunction^[31]^. Accordingly, our findings suggest that DRG neurons in SOD1G93A mice exhibit early molecular changes that resemble those observed in neuropathic disorders, even at the onset stage of ALS.

Electrophysiological recordings revealed enhanced membrane excitability in A-fiber DRG neurons of ALS model mice at disease onset. The proportion of neurons exhibiting repetitive firing also tended to increase in the ALS group, suggesting a functionally hyperexcitable state. Increased action potential amplitude suggests augmentation of inward Na⁺ currents during the rising phase of the action potential, consistent with altered voltage-gated sodium channel function. In addition, depolarization of the RMP and a reduction in rheobase indicate that action potentials can be elicited by smaller depolarizing inputs, further supporting heightened excitability in A-fiber neurons. Because A-fiber DRG neurons mediate tactile and proprioceptive signaling, these electrophysiological changes may provide a mechanistic basis for sensory disturbances, including numbness and dysesthesia, reported in a subset of patients with ALS^6^.MN hyperexcitability in early ALS has been attributed in part to increased persistent Na⁺ currents mediated by Nav1.6^[17]^. In contrast, we did not detect significant changes in Nav1.6 colocalization in DRG neurons. Instead, increased localization of Nav1.7 and Nav1.8 was observed, aligning with the enhanced excitability detected electrophysiologically. These findings suggest that although oxidative stress may represent a shared upstream driver of hyperexcitability in both motor and sensory neurons at disease onset, the downstream molecular mechanisms may differ. Nav1.6 may predominate in MNs, whereas Nav1.7 and Nav1.8 appear more central to excitability changes in DRG neurons. Collectively, the data support the concept that early-stage ALS involves hyperexcitability in primary sensory neurons in addition to motor neurons.

In the present study, increased colocalization of Nav1.7 and Nav1.8 was observed in DRG neurons of ALS model mice at disease onset, with the most pronounced changes in A-fiber neurons. Nav1.7 plays a critical role in setting action potential threshold, whereas Nav1.8 contributes to persistent Na⁺ currents and repetitive firing^[9,10]^. The observed redistribution of these channels is therefore consistent with the enhanced membrane excitability detected electrophysiologically in A-fiber DRG neurons. Altered expression or function of Nav1.7 and Nav1.8 has been widely reported in pain and neuropathy models^[11,12,26,32]^, and oxidative stress is known to modulate both channel expression and function of these channels^[11,12,29]^. Oxidative stress can induce structural changes and altered gating properties in Nav channels, thereby promoting neuronal hyperexcitability^[11]^. In diabetic peripheral neuropathy, increased oxidative stress is associated with upregulation of Nav1.7 and Nav1.8, RMP depolarization, and reduced rheobase^[29,33–37]^—changes that closely resemble the excitability phenotype observed here.

These parallels suggest that Nav channel alterations may represent a common downstream mechanism linking oxidative stress to sensory neuron dysfunction. A fibers, which are large-diameter and fast-conducting, have high metabolic demands and are considered particularly vulnerable to oxidative stress^[18]^. This vulnerability is consistent with the fiber-type–specific Nav channel changes and hyperexcitability observed in A fibers but not in C fibers. Given that sensory disturbances and small-fiber pathology have been described in both patients with ALS and experimental models^[6,23,39,40]^, the A-fiber Nav channel alterations identified here may contribute to the mechanisms underlying sensory abnormalities in ALS.

This study has several limitations. First, the RNA-seq analysis was performed on bulk DRG tissue and therefore could not resolve cell-type–specific transcriptional changes among sensory neurons, satellite glial cells, and infiltrating immune cells. Consequently, the precise cellular sources of the observed gene expression alterations remain undefined. Second, although associations were identified among increased *Fos* and *Npy* expression, altered Nav channel localization, and enhanced membrane excitability, causality was not directly tested. Interventional approaches were not performed to determine whether these molecular changes mechanistically drive the electrophysiological phenotype. Third, the study was cross-sectional and limited to disease onset, precluding assessment of temporal dynamics across presymptomatic and progressive stages.

Future investigations should incorporate single-cell RNA-seq to delineate cell-type–specific molecular alterations within the DRG. Genetic or pharmacological manipulation of *Fos*, *Npy*, and Nav channels would enable direct evaluation of their contributions to sensory neuron hyperexcitability. In addition, longitudinal studies spanning presymptomatic through late disease stages will be essential to determine when sensory dysfunction emerges and how it evolves during ALS progression. Such studies may further support the concept that ALS represents a multisystem neurodegenerative disorder extending beyond MNs and may inform the development of therapeutic strategies targeting sensory system involvement.

## Conclusion

This study demonstrated that, in the DRG of ALS model mice at disease onset, coordinated molecular, structural, and functional alterations—including Nav channel alterations, neuronal soma atrophy, and increased membrane excitability—occur on the background of activation of oxidative stress–related pathways. These findings reinforce the concept that ALS extends beyond a motor neuron disorder and provide important insights into the pathophysiological basis of sensory dysfunction, highlighting potential therapeutic targets within the sensory system.

## Materials and Methods

### Ethics statement

All animal experiments were conducted in accordance with the Guidelines for Proper Conduct of Animal Experiments (2006) established by the Science Council of Japan. Experimental protocols were approved by the Osaka University Recombinant DNA Experiment Safety Committee (approval number: 05365) and the Animal Care and Use Committee of the Graduate School of Dentistry, Osaka University (approval number: R-03-010-0).

### Animals

In this study, mice were generated by breeding male ALS model mice (JAX strain: C57B6SJL-Tg) (SOD1G93A)1Gur/J; mSOD1 mice) (Jackson Laboratory, Bar Harbor, MA, USA) with female wild-type (WT) C57BL/6JJmsSlc mice (WT mice)^[16,41]^. Twelve-week-old male mice were used for all experiments. For RNA sequencing and immunofluorescence analyses, WT (N = 3) and mSOD1 (N = 3) mice were included in each group. For electrophysiological recordings, WT (N = 6) and mSOD1 (N = 5) mice were analyzed (Table 7). N represents the znumber of mice, and n represents the number of samples.

**Table 7.** Number of mice used in each experiment.

| mouse | C57BL/6JmsSlc | B6SJL-Tg<br>(SOD1G93A)1Gur/J |
| --- | --- | --- |
| Gene expression analysis | 3 | 3 |
| Immunofluorescence | 3 | 3 |
| Electrophysiology | 6 | 5 |

Mice were housed individually under controlled environmental conditions (23 °C, 60% humidity, 12-hour light/dark cycle) with ad libitum access to standard laboratory chow (MF; Oriental Yeast, Tokyo, Japan) and water^[42]^. In this model, limb tremors and spasms typically appear at approximately 12 weeks of age, and mice reach the humane endpoint at approximately 20 weeks^[43,38]^. Therefore, 12 weeks of age was defined as disease onset for the present study.

### Genotyping

Genomic DNA was extracted from 5-millimeter tail biopsies using the Quick-DNA Miniprep Plus Kit (ZYMO Research, Irvine, CA, USA). Genotyping was performed using polymerase chain reaction (PCR) according to the protocol provided by The Jackson Laboratory (Protocol 29082: Standard PCR Assay – Tg(SOD)) (Table 8) ^[44]^.

**Table 8.**
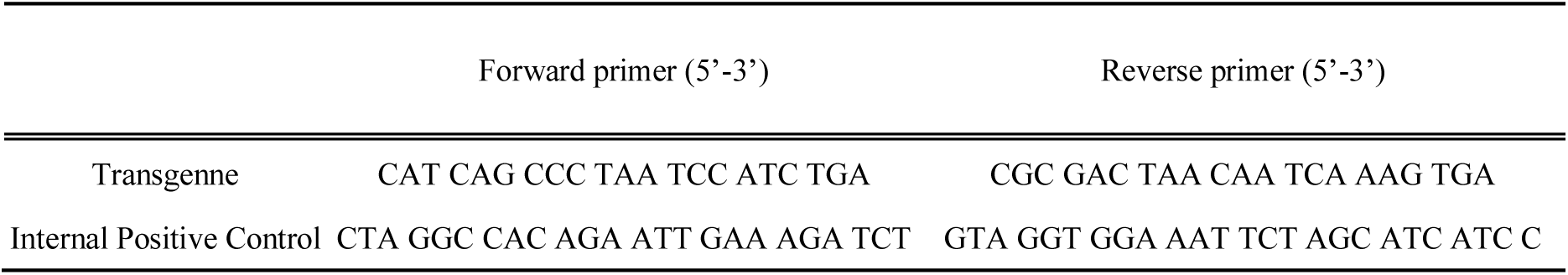
Primers used for genotyping.

### RNA sequencing and bioinformatics

Lumbar DRG were rapidly dissected and immediately frozen in liquid nitrogen. Total RNA was extracted, and cDNA libraries were prepared using the SMART-Seq HT Kit (Takara Bio, Shiga, Japan). Sequencing libraries were subsequently generated using the Nextera XT DNA Library Preparation Kit (Illumina, San Diego, CA, USA) according to the manufacturer’s instructions. High-throughput sequencing was performed on a NovaSeq 6000 platform (Illumina).

Raw FASTQ files were quality-checked using FastQC (v0.12.1) and trimmed using Trimmomatic (v0.33). Cleaned reads were aligned to the *Mus musculus* reference genome (GRCm39; GCF_000001635.27) using STAR (v2.7.0a). Gene-level read counts were generated using featureCounts (v1.5.2)^[45]^.

Differential gene expression analysis was performed using iDEP (v2.4.3)^[46]^. Genes with a false discovery rate (FDR) < 0.05 and an absolute fold change > 2 were considered differentially expressed^[47]^. Functional enrichment analysis was conducted using Gene Set Enrichment Analysis (GSEA) and Kyoto Encyclopedia of Genes and Genomes (KEGG) pathway analysis^[48]^.

Principal component analysis (PCA), heatmaps, and volcano plots were generated using iDEP and the R package ggplot2. Publicly available spinal motor neuron transcriptomic datasets (GSE184484 and GSE281064) were obtained from the Gene Expression Omnibus for comparative analyses^[49,50]^. Some computational analyses were performed using the SQUID high-performance computing system at the D3 Center, Osaka University.

The RNA-seq datasets generated in this study are being deposited in the Gene Expression Omnibus under BioProject accession number PRJNA1419325 (SRA: SRR37134852– SRR37134857; GSE318631).

### Immunofluorescence and imaging

Mice were initially sedated using an intraperitoneal injection comprising medetomidine hydrochloride (0.02 mg/kg), midazolam (0.3 mg/kg), and butorphanol tartrate (0.2 mg/kg). Mice were deeply anesthetized and transcardially perfused with phosphate-buffered saline (PBS), followed by 4% paraformaldehyde (PFA)^[51]^. DRG tissues were dissected, post-fixed for 2 h in 4% PFA, cryoprotected in 30% sucrose, embedded, and sectioned at 5 μm thickness. Sections were then permeabilized and blocked in PBS containing 0.1% Triton X-100 and 5% normal donkey serum, followed by overnight incubation at 4 °C with primary antibodies against NF200 and peripherin (1:1000) and Nav1.6, Nav1.7, and Nav1.8 (1:200) (Table 9). After washing, sections were incubated with Alexa Fluor 488– or Alexa Fluor Plus 555– conjugated secondary antibodies (1:500) for 1 h at 25 °C. Nuclei were counterstained with DAPI^[52]^. Fluorescence images were acquired using a BZ-X800 fluorescence microscope (KEYENCE, Osaka, Japan) under identical exposure settings for all groups.

**Table 9.**
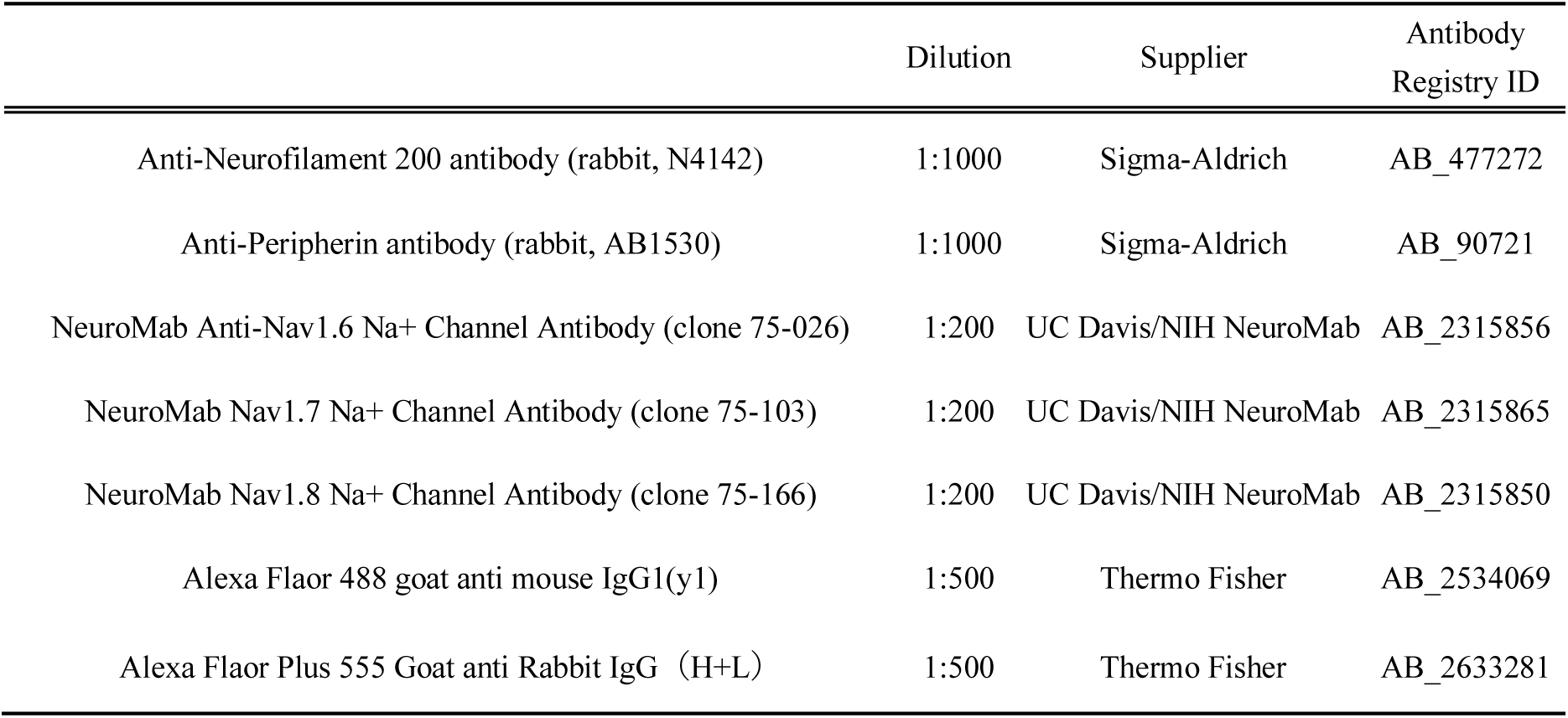
Antibodies used and their dilutions.

### Image quantification

Fluorescence quantification was performed using the ImageJ software^[53]^. Regions of interest (ROIs) corresponding to neuronal somata were manually defined. Mean fluorescence intensity was measured after background subtraction. Nav channel expression levels were quantified as normalized fluorescence intensity ratios.

### Primary DRG neuron culture

Freshly isolated DRG tissues were enzymatically dissociated using collagenase and trypsin to obtain single-cell suspensions^[54]^. Cells were plated onto coverslips coated with poly-L-lysine and laminin and maintained in culture for 24 h at 37 °C in a humidified atmosphere containing 5% CO₂ before electrophysiological recording.

### Electrophysiology

Coverslips containing cultured DRG neurons were transferred to a 2.0 mL acrylic recording chamber mounted on an upright Nomarski infrared differential interference contrast microscope (BX51W1; Olympus). Neurons were continuously perfused with artificial cerebrospinal fluid (ACSF, comprising (n mM) 124 NaCl, 3 KCl, 1.25 NaH₂PO₄, 26 NaHCO₃, 10 glucose, 2 CaCl₂, 2 MgCl₂) at 2 mL/min. Under infrared visualization, DRG neurons were identified as pseudounipolar, round-to-oval cells with diameters of approximately 20–30 µm. Whole-cell patch-clamp recordings were obtained following giga-seal formation. Recording electrodes (3–5 MΩ) were fabricated from borosilicate glass capillaries (outer diameter 1.5 mm, inner diameter 1.12 mm; Intermedical, Tokyo, Japan) using a micropipette puller (P-87; Sutter Instruments, Novato, CA, USA). The internal pipette solution contained (in mM): 115 K-gluconate, 25 KCl, 9 NaCl, 10 HEPES, 0.2 EGTA, 1 MgCl₂, 3 K₂-ATP, and 1 Na-GTP (pH 7.25). Cells with an access resistance <15 MΩ were included in the analysis, and electrical activity was recorded under voltage-clamp (v-clamp) or current-clamp (c-clamp) conditions. Signals were amplified using a Multiclamp 700B amplifier, digitized with an analog-to-digital converter (Digidata 1550A), and acquired using CLAMPEX 10.6 software (all from Molecular Devices, San Jose, CA, USA).

Basic membrane properties were assessed as follows. Membrane input resistance (Rin) was calculated under voltage-clamp conditions using a 5 mV step (10 ms) from a holding potential of −70 mV. Membrane capacitance (Cm) was determined by integrating the capacitive current evoked by a 15 ms hyperpolarizing step. Under current-clamp conditions, resting membrane potential (RMP) was recorded, and action potentials (APs) were elicited by injecting depolarizing current pulses (10 pA, 10 ms). Spike amplitude, afterhyperpolarization (AHP) amplitude, and AHP half-duration were measured. Repetitive firing was induced by 0.5 s step current injections ranging from −500 to 600 pA in 100 pA increments. The presence or absence of firing, the minimum current required to evoke repetitive firing (rheobase), and the mean firing frequency (Hz)—calculated by dividing the number of spikes by time—were recorded. Frequency–current (F–I) curves were generated from the obtained firing frequencies and injected currents^[12,55]^.

## Statistical analysis

Statistical analyses were performed using SPSS software. Data were tested for normality prior to analysis. For comparisons between two groups, Welch’s t-test was used for normally distributed data with unequal variances, and the Mann–Whitney U test was used for nonparametric data. Categorical variables were analyzed using the χ² test. F–I curves were analyzed using two-way repeated-measures analysis of variance (ANOVA), followed by Tukey–Kramer post hoc testing where appropriate. A *p*-value < 0.05 was considered statistically significant. In this study, N denotes the number of animals, and n denotes the number of DRG neurons analyzed.

## Acknowledgements

We would like to thank Editage (www.editage.jp) for English language editing.

## Author Contributions

Conceptualization, S.S.; methodology, A.N., T.K. and S.S.; validation, A.N., T.K., Y.K., S.K., T.Y., Y.O., M, O and S.S.; investigation, A.N., M, O. and S.S.; data collection, A.N., writing original draft preparation, A.N. and S.S.; wright review and editing, S.S.; supervision., S.S. K.M., U.T., T,I., T,Y, Y.Y., E.T.I and S.T.; project administration, S.S. and S.T.; funding acquisition, S.S. All authors have read and agreed to the published version of the manuscript.

## Data availability statement

Raw data were generated at Department of Oral and Maxillofacial Surgery, Graduate School of Dentistry, The University Osaka. Derived data supporting the findings of this study are available from the corresponding author S.S on request.

The datasets generated and/or analysed during the current study are available in the Gene Expression Omnibus (GEO) repository under accession number GSE318631 (https://www.ncbi.nlm.nih.gov/geo/query/acc.cgi?acc=GSE318631). The related BioProject and SRA data are also available under the accession numbers PRJNA1419325 and SRR37134852–857, respectively.

